# Reactive nitrogen species inhibit branched chain alpha-ketoacid dehydrogenase complex and impact muscle cell metabolism

**DOI:** 10.1101/2023.07.31.551364

**Authors:** Nicholas L. Arp, Gretchen Seim, Jordyn Josephson, Jing Fan

## Abstract

Branched chain α-ketoacid dehydrogenase complex (BCKDC) is the rate limiting enzyme in branched chain amino acid (BCAA) catabolism, a metabolic pathway with great importance for human health. BCKDC belongs to the mitochondrial α-ketoacid dehydrogenase complex family, which also includes pyruvate dehydrogenase complex (PDHC) and oxoglutarate dehydrogenase complex (OGDC). Here we revealed that BCKDC can be substantially inhibited by reactive nitrogen species (RNS) via a mechanism similar to what we recently discovered with PDHC and OGDC — modifying the lipoic arm on its E2 subunit. In addition, we showed that such reaction between RNS and the lipoic arm of the E2 subunit can further promote inhibition of the E3 subunits of α-ketoacid dehydrogenase complexes. We examined the impacts of this RNS-mediated BCKDC inhibition in muscle cells, an important site of BCAA metabolism, and demonstrated that the nitric oxide production induced by cytokine stimulation leads to a strong inhibition of BCKDC activity and BCAA oxidation in myotubes and myoblasts. More broadly, nitric oxide production reduced the level of functional lipoic arms across the multiple α-ketoacid dehydrogenases and led to intracellular accumulation of their substrates (α-ketoacids), reduction of their products (acyl-CoAs), and a lower cellular energy charge. This work revealed a new mechanism for BCKDC regulation, demonstrated its biological significance, and elucidated the mechanistic connection between RNS-driven inhibitory modifications on the E2 and E3 subunits of α-ketoacid dehydrogenases. Together with previous work, we revealed a general mechanism for RNS to inhibit all α-ketoacid dehydrogenases, which has numerous physiological implications across multiple cell types.

## Introduction

The family of α-ketoacid dehydrogenase complexes, which includes pyruvate dehydrogenase complex (PDHC), oxoglutarate dehydrogenase complex (OGDC), and branched chain α-ketoacid dehydrogenase complex (BCKDC), play key roles in mitochondrial metabolism. These enzyme complexes share a similar catalytic mechanism involving coupled action of three subunits (Fig 1*A*). The E1 subunit, PDH, OGDH, and BCKDH for the three enzyme complexes, respectively, is a thiamin-dependent decarboxylase that decarboxylates its substrate (α-ketoacids) to its corresponding acyl-group. The E2 subunits, dihydrolipoamide S-acetyltransferase (DLAT), dihydrolipoamide S-succinyltransferase (DLST), and dihydrolipoamide branched chain transacylase (DBT), respectively, contain a covalently attached lipoic arm that mediates the transfer of the acyl-group from the E1 subunit to coenzyme A (CoA) to produce acyl-CoA. With this transfer, the lipoic arm converts from its oxidized form (lipoamide) to its reduced form (dihydrolipoamide). Finally, the E3 subunit, dihydrolipoamide dehydrogenase (DLD), encoded by the same gene for all three α-ketoacid dehydrogenase complexes, re-oxidizes the dihydrolipoamide to lipoamide, coupled to NAD reduction to NADH. Together, it allows for the oxidation of α-ketoacids and the production of acyl-CoA and NADH. These enzymes catalyze reactions that are key steps in carbohydrates and amino acids catabolism (1).

**Figure 1.**
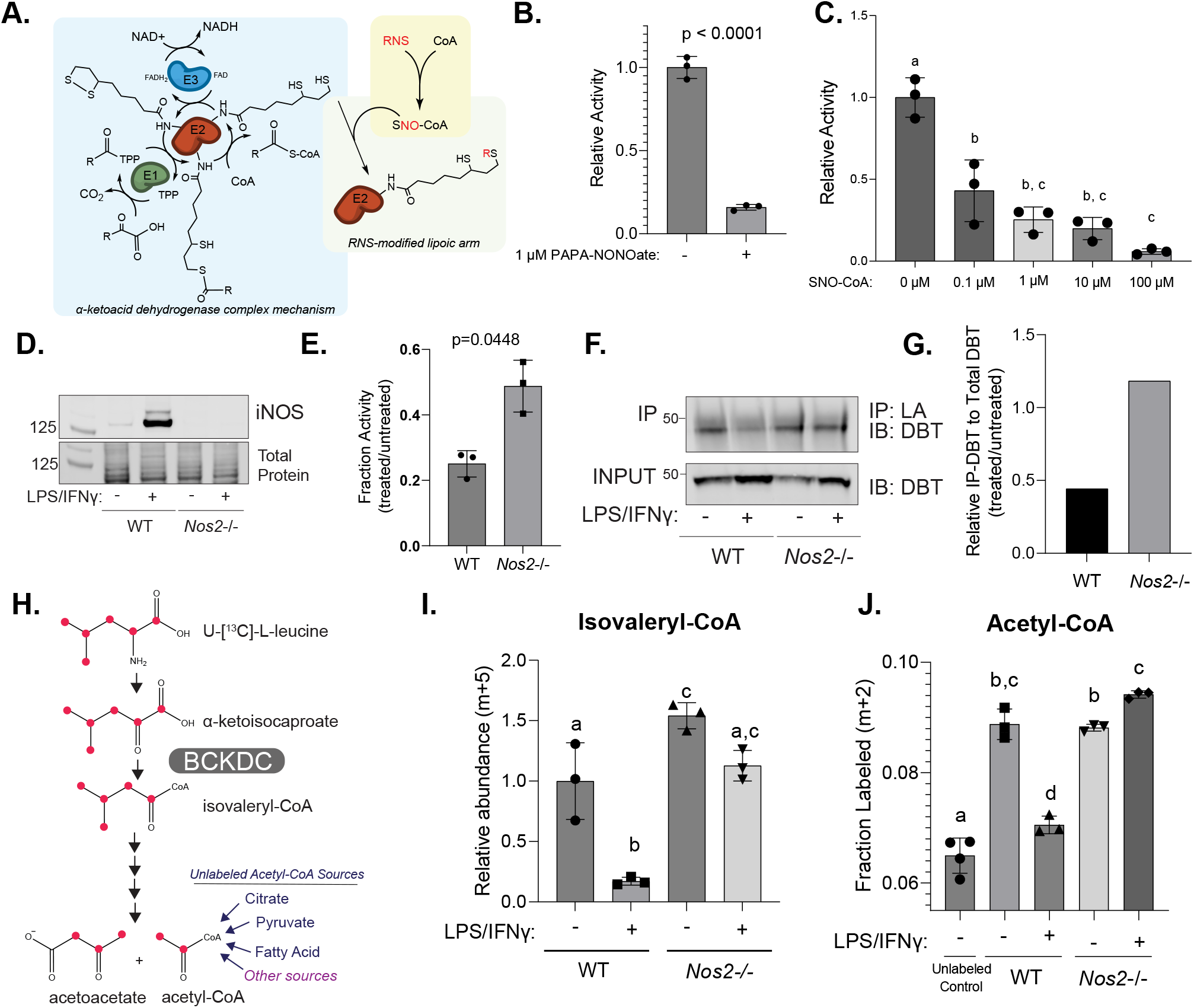
**A.** *Model schematic.* RNS can cause inhibitory modification of the E2 subunit’s catalytic lipoic arm, through the targeted delivery via SNO-CoA. **B.** Relative activity of BCKDC in mitochondria lysate from wild type RAW264.7 cells, after *in vitro* incubation with PAPA-NONOate (1 mM) for 5 hours at room temperature. Statistical analysis was performed with unpaired student’s t-test with p-value reported. **C.** Relative BCKDC activity in matrix lysate from wild type RAW264.7 cells after *in vitro* incubation with varied doses of SNO-CoA in the presence of NADH (200 µM) for 3 hours at room temperature. Statistical analysis was performed using one-way ANOVA followed by Tukey’s post-hoc test. Columns with different letters indicate a statistical significance of p<0.05. **D.** Immunoblot for iNOS in whole cell lysate of wild type (WT) and *Nos2*−/− RAW264.7 cells with or without LPS and IFNγ stimulation. **E.** Relative BCKDC activity in LPS and IFNγ activated state (24-hour post stimulation) compared to unstimulated state in wild type (WT) or *Nos2*−/− RAW264.7 cells. Statistical analysis was performed using unpaired student’s t-test with exact p-value reported. **F-G**. the level of functional lipoic arm on DBT in wild type (WT) or *Nos2*−/− RAW264.7 cells in unstimulated state or 24 hours after LPS and IFNγ stimulation. (F) immunoblots of DBT in RAW264.7 whole cell lysates (input) or after Immunoprecipitation of lipoic acid. (G) relative fraction of DBT with functional lipoic arm in stimulated state compared to unstimulated state, quantified based on blots in (*F)*. **H.** U-[^13^C]-L-leucine isotopic tracing schematic. Red dots represent ^13^C conversion through the leucine catabolic pathway. **I.** Relative abundance of isovaleryl-CoA (M+5 labeled from U-[^13^C]-L-leucine) in wild type (WT) or *Nos2*−/− RAW264.7 cells with or without LPS and IFNγ stimulation for 24 hours. **J.** The fraction of M+2 labeled acetyl-CoA after 48 hours labeling in media containing U-[^13^C]-L-leucine (n=3) or unlabeled control (n=4) in wild type (WT) or *Nos2*−/− RAW264.7 cells with or without LPS and IFNγ stimulation for 24 hours. **I and J.** Statistical analysis was performed using one-way ANOVA followed by post-hoc Tukey’s test. Columns with different letters indicate a statistical significance of p<0.05. **B, C, E, I, and J**. bars and error bars represent mean ± standard deviation, n = 3 unless otherwise noted.

Specifically, BCKDC is the rate liming step in the oxidation of the branched chain amino acids (BCAA), valine, isoleucine, and leucine. The catabolism of these essential amino acids contributes carbon units to various metabolic pathways, including the tricarboxylic acid (TCA) cycle, ketone body metabolism, and lipid synthesis. In addition, BCAAs and their catabolic intermediates play important roles in nutrient sensing and signaling (2, 3). The first step of BCAA catabolism is mediated by branched chain aminotransferase, Bcat1 or Bcat2, which transfers the amino group from valine, isoleucine, or leucine to alpha-ketoglutarate, producing the corresponding α-ketoacids, α-ketoisovalerate (αKIV), α-keto-β-methylvalerate (αKMV), and α-ketoisocaproate (αKIC), respectively. Next, BCKDC oxidizes these α-ketoacids to their corresponding acyl-CoA products, α-methylbutyryl-CoA, isobutyryl-CoA, and isovaleryl-CoA, respectively. BCAA oxidation flux is particularly high in skeletal muscle (4), where it plays a key role in systemic metabolism and human health (5–7). Dysregulated BCAA metabolism is associated with many diseases, including cardiovascular disease, such as atherosclerosis and heart failure, and metabolic disorders, such as diabetes mellitus and obesity (2, 8–12).

As BCKDC, PDHC, and OGDC control important crossroads of the metabolic network, their activities are dynamically regulated by layers of molecular mechanisms. Particularly, multiple mechanisms acting through post-translational modifications of E1, E2, or E3 subunits of these enzymes have been found to play key roles in their regulation (13–23). Recently, we discovered that reactive nitrogen species (RNS) produced in classically activated macrophages by inducible nitric oxide synthase (iNOS), encoded by the gene *Nos2*, led to profound inhibition of PDHC and OGDC by a previously unknow mechanism (24): This inhibition is caused by profound reduction of functional lipoic arm on their E2 subunit, as can be detected by immunoblotting against lipoic moiety. However, mass spectrometry-based analysis showed this reduction is largely not due to de-lipoylation, rather, RNS can cause a series of covalent S-modifications of the lipoic arm, which prevent the lipoic arm from cycling between its reduced and oxidized forms to perform its catalytic activity (Fig. 1*A*). Furthermore, we demonstrated that the RNS-driven inhibition specifically acts through thiol modification of the E2 subunit’s lipoic arm, by showing inhibition of purified PDHC by RNS depends on the presence of its substrates (pyruvate, CoA, or NADH). Without the addition of substrates, E2 subunit’s lipoic arm is mainly in its oxidized form without reactive thiol. In this condition, NO donor alone does not cause substantial inhibition of PDHC. In contrast, incubating purified PDHC with its substrates, pyruvate and CoA, or its product NADH, allows the lipoic arm to convert to its reduced form via E1 and E2 subunits’ activity or reversed E3 subunit activity respectively, exposing the reactive thiols. Consistently, incubating purified PDHC with NO donor in the presence of pyruvate and CoA, or NADH, causes substantial inactivation of PDHC. Given the mechanistic similarity among α-ketoacid dehydrogenase complexes, it is conceivable that a similar mechanism could also regulate BCKDC. Additionally, many other cell types beyond macrophages also produce nitric oxide (NO), therefore, it is also likely that the RNS-driven regulation of these complexes is has importance in broad contexts. However, these hypotheses have not been tested.

Here through a series of *in vitro* and in cell experiments, we show that RNS can inhibit BCKDC as well via inactivating thiols modification of the lipoic arm on its E2 subunit. This mechanism can not only inhibit BCKDC alongside PDHC and OGDC in classically activated macrophages, but also occurs in muscle cells, in which BCKDC plays key physiological roles. Upon exposure to tumor necrosis factor-alpha (TNF-α) and interferon-gamma (IFN-γ), muscle cells express iNOS and produce NO (25). This NO production has been implicated in muscle cachexia and altered mitochondrial metabolism (26). We found that TNF-α and IFN-γ stimulated NO production in myotubes and myoblasts lead to significant inhibition of BCKDC, PDHC, and OGDC, and altered energy charge.

In addition to targeting E2 subunit’s lipoic arm, RNS have also been shown to inhibit α-ketoacid dehydrogenase complex through mechanisms acting on the E3 subunit (DLD). In classically activated macrophages, RNS can cause inhibitory cysteine nitrosylation of DLD (21). The normal catalytic function of DLD in α-ketoacid dehydrogenase complexes is to re-oxidize the reduced lipoic arm on E2 subunit using a cysteine-cysteine active site. Based on this close interaction between the subunits, we reasoned that the RNS-driven modifications on the E2 subunit’s lipoic arm could promote the modification of E3 subunits. Indeed, here we show that the RNS-driven inhibition of the E3 subunit largely depends on lipoic arm modification of the E2 subunit, providing a mechanistic link between the two recently discovered RNS-driven inhibitory mechanisms.

Together, these data demonstrate a common mechanism which allows RNS to inhibit important enzymes across the lipoic arm-dependent dehydrogenase family, including BCKDC. Such regulation by RNS has significant biological impacts in multiple cell types that are capable of producing RNS and is likely to have broader relevance in other cell types that are influenced by RNS in the microenvironment under specific physiological and pathological conditions.

## Results

### RNS cause strong inhibition of BCKDC

To test the hypothesis that NO can inhibit BCKDC, we first incubated mitochondrial lysate from the macrophage cell line, RAW 264.7 cells, with NO donor, PAPA-NONOate. Indeed, *in vitro* treatment with NO donor led to a profound inhibition of BCKDC activity (Fig. 1*B*).

We hypothesized this inhibition was mediated mainly through a mechanism similar to what we recently found with PDHC and OGDC (24): RNS cause a series of inactivating S-modifications of the lipoic arm on their E2 subunit, and such mechanism can be highly specific and efficient, because RNS can react with cellular CoA to form SNO-CoA, which, via binding to their E2 subunit at the CoA binding site, delivers the modifications to the lipoic arm in a targeted manner. To test if BCKDC can be directly inhibited by SNO-CoA, we incubated mitochondrial lysate with varied doses of SNO-CoA in the presence of NADH (to generate reduced thiol on lipoic arm) and measured BCKDC activity. Indeed, SNO-CoA inhibited BCKDC in a dose dependent manner, with concentration as low as 0.1μM can cause over 50% activity reduction in 3 hours, and 10μM near completely inactivated BCKDC (Fig. 1*C*). These results provided *in vitro* evidence that BCKDC can be efficiently inhibited by RNS similar to PDHC and OGDC.

### NO production leads to BCKDC inhibition in activated macrophages

We next tested the hypothesis that such RNS-driven inhibition of BCKDC occurs in cells. In macrophages, classical activation by lipopolysaccharide (LPS) and interferon-γ (IFNγ) induces the expression of iNOS (Fig. 1*D*), resulting in the production of NO. In the macrophage cell line, RAW 264.7, BCKDC activity is reduced by ∼80% upon classical activation, and such BCKDC inhibition upon activation is significantly rescued by *Nos2* knock out (Fig. 1*E*), showing NO is an important driver of the BCKDC inhibition in activated macrophages. To test whether this NO-dependent inhibition of BCKDC is mediated by changes in the lipoic arm, we probed for the level of functional lipoic arm on BCKDC’s E2 subunit, DBT, by immunoprecipitation. Although BCKDC activity decreased substantially upon activation, the total level of DBT is slightly higher upon activation in both wildtype and *Nos2*−/− macrophages (Fig. 1*F*), possibly due to compensation, suggesting the activation-induced inhibition is not due to reduced DBT level, but strong inactivation of its catalytic activity. Correlating with overall BCKDC activity, the level of functional lipoic arm on DBT decreased substantially upon activation in wild type macrophages, but such decrease was prevented by *Nos2* knock out (Fig. 1, *E, F* and *G*), suggesting that NO causes changes to the lipoic arm which then mediate the BCKDC inhibition upon activation.

BCKDC is the rate limiting step in BCAA catabolism. To examine the impact of NO production on BCAA metabolism in macrophages, we applied isotopic tracing with U-[^13^C]-L-leucine. Oxidation of U-[^13^C]-L-leucine by BCKDC produces 5-labeled isovaleryl-CoA, which can be further metabolized to labeled acetyl-CoA, whereas the unlabeled fraction of acetyl-CoA originates from other sources, including citrate, pyruvate, and β-oxidation of fatty acids (Fig. 1*H*). In wild type RAW264.7 cells, stimulation by LPS and IFNγ greatly reduced the abundance of 5-labeled isovaleryl-CoA and reduced the contribution of U-[^13^C]-L-leucine to acetyl-CoA production to near-blank level (Here the blank was the fraction of M+2 acetyl-CoA measured in cells cultured in fully unlabeled media. The M+2 arises from natural abundance of C13 and S34). Both the stimulation-induced reduction in isovaleryl-CoA abundance and in acetyl-CoA labeling from leucine were rescued by *Nos2* knock out (Fig. 1, *I* and *J*). Together, these results show NO production causes lipoic arm alteration and inhibition of BCKDC, and reduction in BCAA oxidation, in macrophages upon classical activation.

### NO inhibits α-ketoacid dehydrogenase complexes in myotubes and myoblasts

Many tissues and cell types have the capability of producing NO by nitric oxide synthases for functions including signaling, pathogen killing, and regulation of angiogenesis, vasodilation, and neural functions (27–30). Additionally, cells that do not actively produce NO themselves can be impacted by RNS in the microenvironment generated by neighboring cells. Therefore, this mechanism for RNS to inhibit of α-ketoacid dehydrogenase complexes is likely to have broad biological significance in a variety of cellular systems beyond macrophages. Skeletal muscle is known to express iNOS and produce NO upon stimulation with the cytokines, TNF-α and IFN-γ (25). And skeletal muscle is an important site for BCAA catabolism (4). We hypothesized that α-ketoacid dehydrogenase complexes are inhibited by RNS in muscle cells upon cytokine stimulation and tested this hypothesis using a widely used cell model, C2C12 myoblast cells, which can be differentiated to myotubes (31–33).

In differentiated C2C12 myotubes stimulated with TNF-α and IFN-γ for 48 hours, iNOS expression is induced, and the levels of functional lipoic arm in both the bands corresponding to the molecular weight of DLAT (E2 subunit of PDHC, ∼70kDa) and DLST or DBT (E2 subunit of OGDC and BCKDC, respectively, both ∼ 50 kDa) are significantly reduced relative to total level of their corresponding E2 subunits (Fig. 2*A*). Treating cells with a selective inhibitor of iNOS, N-(3-(aminomethyl) benzyl)-acetamidine (1400W), rescued the stimulation-induced decrease in the functional lipoic arm relative to E2 subunit level (Fig. 2*A*). Consistent with the changes in lipoic arm, BCKDC activity, as measured in isolated mitochondria lysate, is reduced by ∼50% upon stimulation, and this inhibition is rescued by iNOS inhibition (Fig. 2*B*).

**Figure 2.**
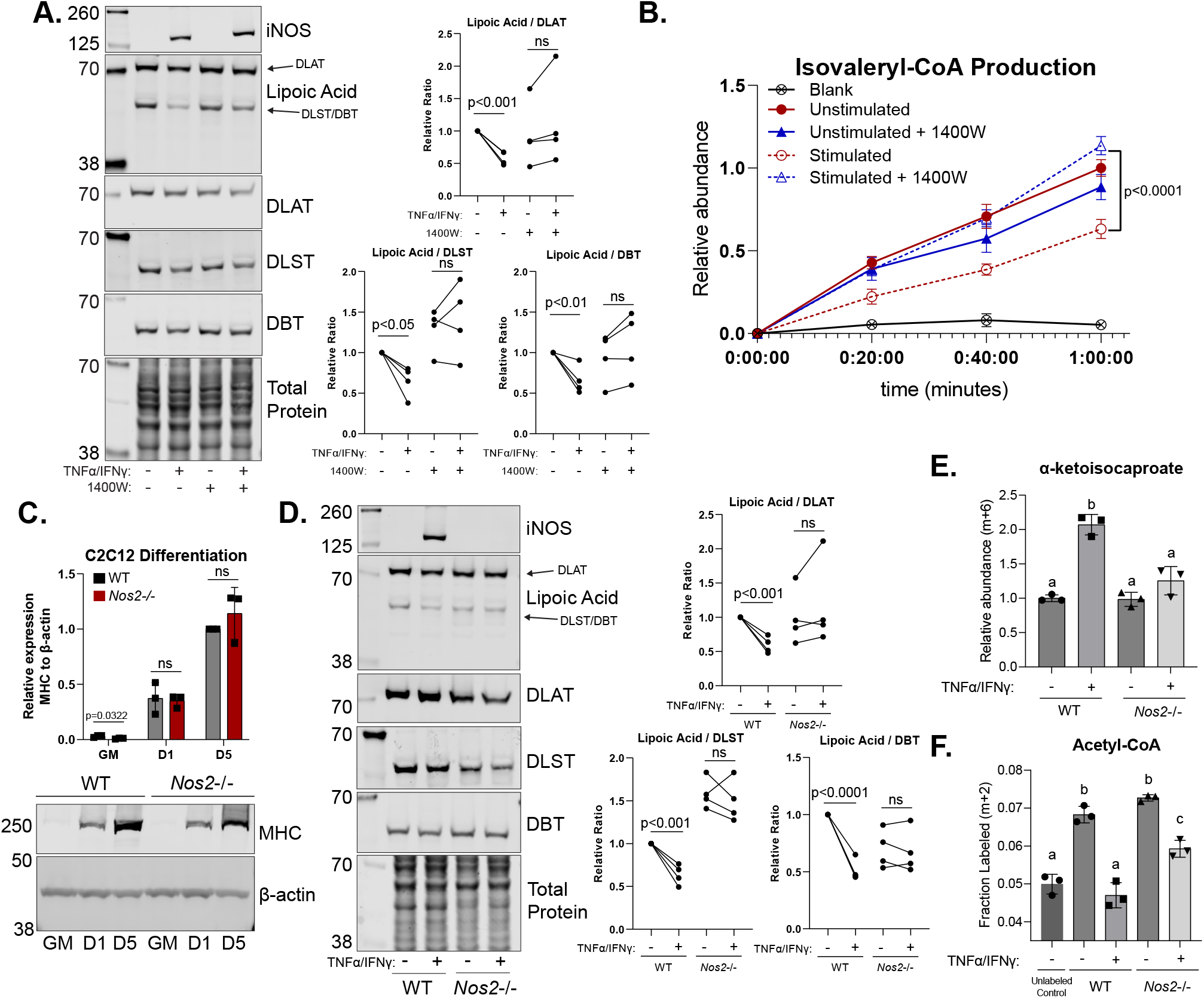
**A.** Representative immunoblots for iNOS, DLAT, DLST, DBT, and lipoic moiety in whole cell lysates of unstimulated C2C12 myotubes or myotubes stimulated with TNFα and IFNγ for 48 hours, with or without treatment with 1400W (200 µM). The experiment was repeated four times. To compare the changes in functional lipoic arm upon stimulation, the relative ratio of lipoic band to its corresponding total E2 subunit band (DLAT for the lipoic band at ∼70kDa, and DLST or DBT for the lipoic band at ∼50kDa) was quantified and normalized to unstimulated, untreated condition, as graphed in the corresponding figure. Each dot represent result for each independent experiments **B**. BCKDC activity, as measured by isovaleryl-CoA production from αKIC over time in crude mitochondria isolation from C2C12 myotubes treated with or without 1400W (200 µM) and with or without TNFα and IFNγ stimulation. **C.** Representative immunoblot for myosin heavy chain (MHC) and beta-actin over C2C12 differentiation time course from wild type and *Nos2−/−* whole cell lysate when cells were in growth media (GM) and after the first (D1) and fifth day (D5) in differentiation media. This experiment was repeated three times. MHC expression was quantified and normalized to loading control (beta-actin) in each experiment. Relative expression was graphed as relative level compared to wild type D5 condition. Each dot represents an independent experiment. **D.** The same as Fig. 2A but with genetic *Nos2* knock out instead of 1400W treatment. **E.** Relative abundance of αKIC (6-labeled from U-[^13^C]-L-leucine) in stimulated (TNFα and IFNγ for 48 hours) or unstimulated wild-type or *Nos2*−/− C2C12 myotubes. **F.** The fraction of M+2 labeled acetyl-CoA after 48 hours labeling with U-[^13^C]-L-leucine in stimulated or unstimulated wild-type or *Nos2*−/− C2C12 myotubes. **A, C, and D.** For immunoblot quantification, statistical analysis was performed with unpaired student’s t-test with p-value of significant changes indicated on graph. ns indicate no significant change (p>0.05). **B**. Plots represent mean ± standard deviation, n = 3. Statistical analysis was performed with unpaired student’s t-test with p-value significant indicated on graph. **E and F.** All bars and error bars represent mean ± standard deviation, n = 3. Statistical analysis for significance was performed with one-way ANOVA followed by a post-hoc Tukey’s test. Columns with different letters indicate a statistical significance of p<0.05.

We further tested the effect of NO on the changes in lipoic arm of α-ketoacid dehydrogenase complexes using a genetic knock out of *Nos2*. Like wildtype C2C12 cells, *Nos2* knockout cells can be sufficiently differentiated to myotubes, as indicated by the great increase of myosin expression after differentiation (Fig 2*C*). Similar to what is observed with pharmacological inhibition of iNOS, the stimulation-induced decrease of functional lipoic arm relative to total DLAT, DLST and DBT level was prevented in *Nos2*−/− myotubes (Fig. 2*D*).

To further probe BCKDC activity in cells, we performed U-[^13^C]-L-leucine tracing. Upon stimulation with TNF-α and IFN-γ, cellular level of labeled αKIC, the substrate of BCKDC, accumulated in wild-type myotubes but not *Nos2*−/− myotubes (Fig. 2*E*). The fraction of 2-labeled acetyl-CoA decreased to blank level upon stimulation in wild type myotubes, and this loss of labeling incorporation from leucine to acetyl-CoA is significantly reversed by *Nos2*−/− (Fig. 3*F*). This NO-dependent buildup of substrate and decrease in labeling incorporation into downstream metabolite indicated NO inhibits BCKDC activity in cells.

**Figure 3.**
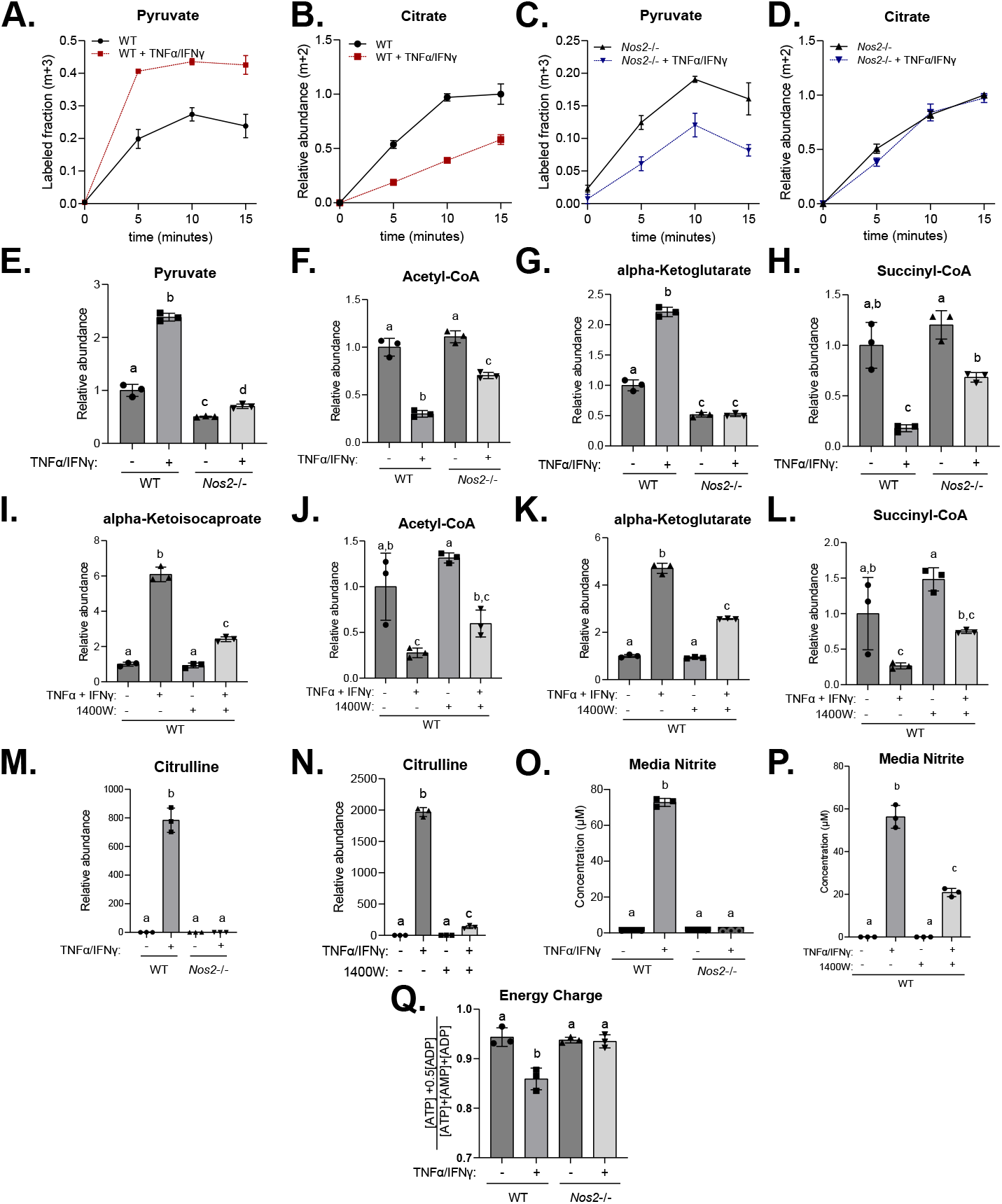
**A and B.** Kinetic labeling incorporation from U-[^13^C]-D-glucose to (A) pyruvate (M+3 labeled fraction) and (B) Citrate (m+2 total abundance) in unstimulated or stimulated (TNFα and IFNγ for 48 hours) wild type C2C12 myotubes. **C and D.** Same as A and B, expect with Nos2−/− C2C12 myotubes. **A-D** Each time point and error bars represent mean ± standard deviation, n = 3. **E, F, G, and H.** Relative abundance of intracellular pyruvate (E), acetyl-CoA (F), α-ketoglutarate (G), and succinyl-CoA (H) in stimulated (TNFα and IFNγ for 48 hours) or unstimulated wild-type or *Nos2*−/− C2C12 myotubes. **I, J, K, and L.** Relative abundance of intracellular αKIC (I), acetyl-CoA (F), α-ketoglutarate (G), and succinyl-CoA (H) in stimulated or unstimulated wild-type C2C12 myotubes with or without 1400W (200 µM). **M and N.** Relative abundance of citrulline in wild type or Nos2−/− C2C12 myotubes (M), or wild type with or without 1400W (200 µM) (N). **O and P.** Nitrite concentration in spent media after culturing wild type or Nos2−/− C2C12 myotubes (O), or wild type with or without 1400W (200 µM) (P). **Q.** Energy charge of stimulated or unstimulated wild-type or *Nos2*−/− C2C12 myotubes. **E-Q.** Statistical analysis was performed with one-way ANOVA followed by a post-hoc Tukey’s test. Columns with different letters indicate a statistical significance of p<0.05. All bars and error bars represent mean ± standard deviation, n = 3.

Based on the observed changes in the lipoic arm (Fig. 2, *A* and *D*), we also expected PDHC and OGDC to be inhibited in a NO-dependent manner in TNF-α and IFN-γ stimulated myotubes. To examine PDHC activity and glucose oxidation in myotubes, we performed kinetic labeling with U-[^13^C]-glucose tracer. U-[^13^C]-glucose is quickly converted to U-[^13^C]-pyruvate via glycolysis in both stimulated and unstimulated condition (Fig. 3*A*). The labeled pyruvate can be metabolized by PDHC to 2-labeled acetyl-CoA, which is then converted to 2-labeled citrate. In wild type myotubes, TNF-α and IFN-γ stimulation caused the labeling incorporation from glucose into citrate to be much slower (Fig. 3*B*), even though the labeling incorporation into pyruvate is higher (Fig. 3*A*), suggesting greatly reduced flux through PDHC. In contrast, this reduced rate of labeling incorporation from pyruvate into citrate was not observed when *Nos2*−/− myotubes were activated (Fig. 3, *C* and *D*). These results suggest NO inhibits intracellular PDHC activity in activated myotubes. Consistently, we observed substantial accumulation of pyruvate, the substrate of PDHC, and depletion of acetyl-CoA, the product of PDHC, in stimulated wild type myotubes, and these stimulation-induced changes are largely prevented by *Nos2*−/− (Fig. 3, *E* and *F*). Similarly, the substrate for OGDC, α-ketoglutarate, accumulated significantly, and the product of OGDC, succinyl-CoA, depleted upon stimulation in wild type myotubes, but these changes are also rescued in *Nos2*−/− myotubes (Fig. 3, *G* and *H*).

Similar to genetic *Nos2* knock out, treating cells with iNOS inhibitor 1400W also partially reversed the stimulation-induced accumulation of α-ketoglutarate and α-ketoisocaproate and depletion of succinyl-CoA and acetyl-CoA (Fig 3, *I, J, K*, and *L*). However, the rescue was weaker as compared to the genetic knock out. This correlates with the fact that genetic *Nos2* knock out completely ablated NO production but 1400W treatment did so incompletely, as evidenced by measurements of intracellular citrulline, a product of iNOS (Fig. 3, *M* and *N*), and extracellular nitrite (Fig. 3, *O* and *P*), suggesting possible dose-dependent effect of NO.

As the results above suggested NO production causes inhibition of all three α-ketoacid dehydrogenase complexes, PDHC, OGDC and BCKDC, which control the mitochondrial oxidation of important nutrients – glucose, glutamine, and BCAA, respectively – such inhibition can have significant impact on cellular bioenergetics. Indeed, consistent with a previous report (26), we observed cellular energy charges significantly decreased upon stimulation of myotubes, and this decrease is prevented by *Nos2*−/− (Fig. 3*Q*).

Like myotubes, myoblasts can also produce NO. Upon stimulation with LPS and IFNγ, C2C12 myoblasts express iNOS (Fig. 4*A*) and accumulate intracellular citrulline (Fig. 4*B*), indicating activated NO production. Consistent with NO driving inhibition of α-ketoacid dehydrogenases, we found similar alterations in stimulated myoblasts, including reduced incorporation of labeled glucose into TCA cycle intermediates (Fig. 4, *C*, *D*, *E*, and *F*), accumulation of α-ketoacids (Fig. 4, *G*, *H*, and *I*), and decrease of their corresponding acyl-CoAs in stimulated myoblasts (Fig. 4, *J* and *K*).

**Figure 4.**
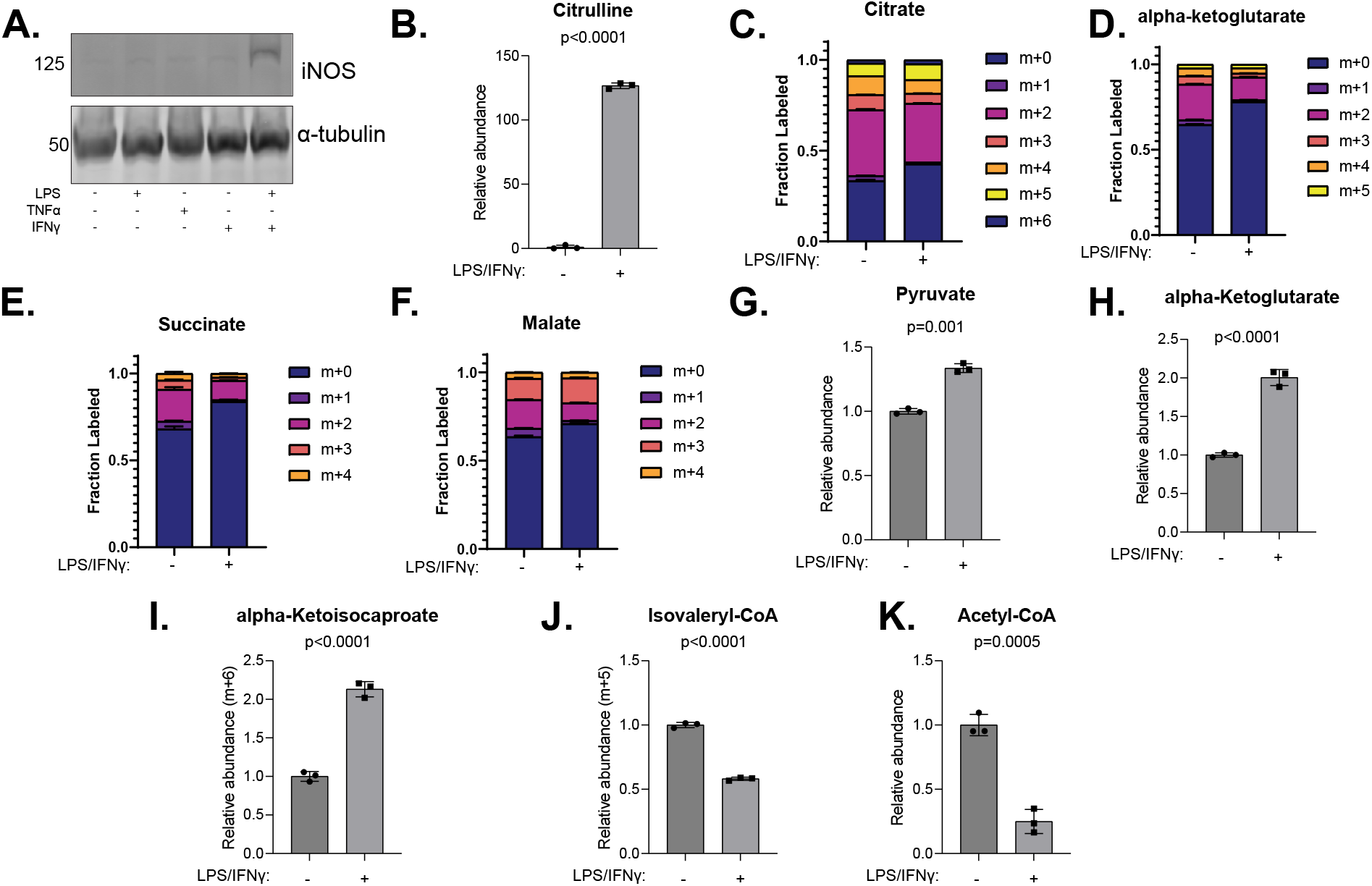
**A.** Immunoblot for iNOS and alpha tubulin in whole cell lysate of C2C12 myoblasts that are unstimulated or stimulated with LPS and IFNγ for 24 hours. **B.** Relative abundance of intracellular citrulline in C2C12 myoblasts with or without 24-hour LPS and IFNγ stimulation. **C, D, E, and F.** Labeling patterns of citrate (C), α-ketoglutarate (D), succinate (E), and malate (F) after unstimulated or LPS and IFNγ stimulated C2C12 myoblasts were cultured in media containing U-[^13^C]-D-glucose for 24 hours. **G and H.** Abundance of pyruvate (G) and α-ketoglutarate (H) in C2C12 myoblasts with or without 24-hour LPS and IFNγ stimulation. **I and J.** Relative abundance of αKIC (6-labeled from U-[13C]-L-leucine) (I) and isovaleryl-CoA (5-labeled from U-[13C]-L-leucine) (J) in C2C12 myoblasts with or without 24-hour LPS and IFNγ stimulation cultured in media containing U-[13C]-L-leucine for 24 hours. **K.** Relative abundance of acetyl-CoA in C2C12 myoblasts with or without stimulation. **B, G -K.** Statistical analysis was performed with unpaired student’s t-test. All bars and error bars represent mean ± standard deviation, n = 3.

### RNS mediated lipoic modification on E2 subunit promotes E3 subunit inhibition

The α-ketoacid dehydrogenase complexes are subjected to the regulation by many mechanisms. Besides the inhibitory modifications of the lipoic arm on E2 subunit, it has been recently discovered that in macrophages RNS can also inhibit PDHC via another mechanism-- cysteine nitrosylation on its E3 subunit (DLD) (21). The normal catalytic function of DLD is to use a cysteine-cysteine active site to re-oxidize the reduced lipoic arm on E2 subunit, then transfer the electron to FAD, then NAD, to produce NADH (Fig. 1*A*). Based on this close interaction between E2 and E3 subunits, we hypothesized that the RNS-driven lipoic modification on E2 subunit can further promote the posttranslational modification on E3 subunit through mechanisms such as trans-nitrosylation.

To test this hypothesis, we took advantage of the fact that, as demonstrated in our previous publication, the modification of E2 subunit’s lipoic arm by RNS depends on the presence of substrates (pyruvate and CoA) to generate reduced thiols that is susceptible to modifications. The presence of CoA is also important for the generation of SNO-CoA to efficiently deliver the modifications to the lipoic arm (Fig. 1*A*) (24). If the modification and inhibition of DLD is resulted from the interaction with E2 subunit with lipoic arm modification, it too would be dependent on the presence of substrates. We therefore incubated purified PDHC with NO donor, PAPA-NONOate, in the presence or absence or pyruvate and CoA and measured the loss of E3 subunit activity. Only when both pyruvate and CoA were present did the NO donor cause a large reduction in overall PDHC activity (Fig. 5*A*) and in specific DLD activity as measured by spectrometric assay (Figure 5*B*) and in-gel activity assay (Fig 5, *C* and *D*) following previously established protocols (21, 34–36). The observation that NO donor can cause substantial inhibition of DLD activity without changing total DLD level in the presence of pyruvate and CoA (Fig. 5, *B*, *C*, and *D*) confirmed that in normal cellular environment, where pyruvate and CoA are present, production of NO could modify and inhibit DLD, as recently reported in macrophages upon classical activation (21). However, the inhibition of DLD activity (18%, Fig 5*B*) is relatively small compared to the inhibition of overall PDHC activity (82%, Fig 5*A*), suggesting the contribution of E3 inhibition is minor and E2 subunit inhibition is the major driver of overall PDHC inhibition by RNS. The fact that without pyruvate or CoA, NO donor alone does not cause significant DLD inhibition demonstrated that NO does not directly cause inhibitory modifications of DLD, and the E3 modification is likely mediated by the interaction with the E2 subunit whose lipoic arm is modified by RNS.

**Figure 5.**
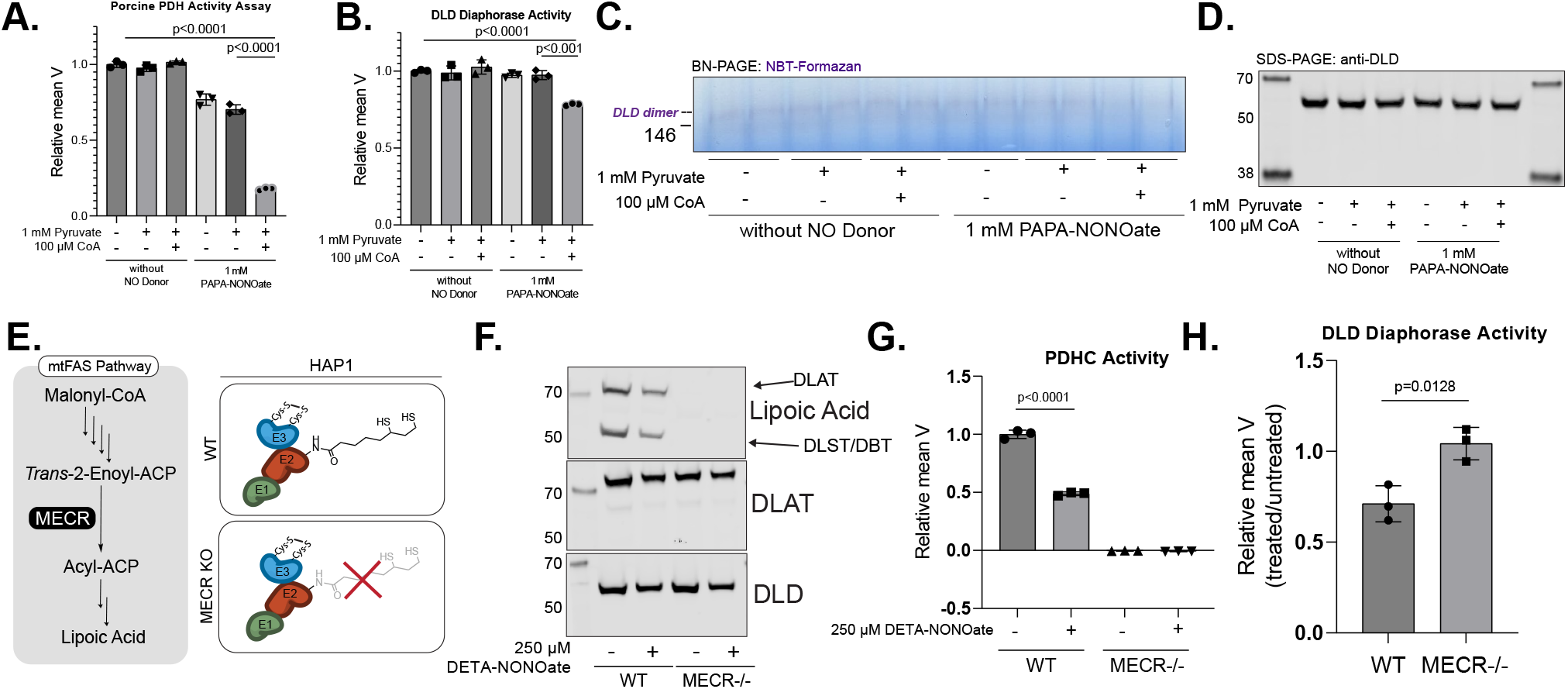
**A.B.** Relative overall PDHC enzymatic activity (A) or DLD activity (B) measured by spectrometric assay after purified porcine PDHC was incubated with indicated combination of PAPA-NONOate (1mM), pyruvate (1mM) and CoA (100 µM). Statistical analysis was performed with student’s t-test, with all significant differences (p<0.05) indicated on figure with the corresponding p-value. All bars and error bars represent mean ± standard deviation, n = 3.**C.** in-gel assay after purified porcine PDHC was incubated with indicated combination of PAPA-NONOate (1mM), pyruvate (1mM) and CoA (100 µM). DLD activity is indicated by purple dye intensity at the molecular weight of DLD dimer (∼146 kDa) in native gel. Samples loaded on gel in duplicates. **D.** Immunoblot for total DLD level in the same samples as (C). **E.** Schematic showing the function of MECR and that MECR knockout HAP1 cells lack lipoic arm on their α-ketoacid dehydrogenase complexes. **F.** Immunoblot for lipoic moiety, DLAT and DLD in wild type and MECR−/− HAP1 cells with or without 250 µM DETA-NONOate treatment. **G.** Relative enzymatic activity of PDHC in wild type or MECR−/− HAP1 cells with or without 250 µ DETA-NONOate treatment. **H.** Relative enzymatic activity of DLD measured by spectrometric assay in DETA-NONOate (250 µM) treated wildtype or MECR−/− cells compared to untreated condition. **G.H.** Statistical analysis was performed with unpaired student’s t-test with p value reported. Bars and error bars represent mean ± standard deviation, n = 3.

To further test this hypothesis in cells, we used a cell model that lacks lipoic arm on α-ketoacid dehydrogenase complexes’ E2 subunit. Mitochondrial trans-2-enoyl-CoA reductase (MECR), a required enzyme in the lipoic acid biosynthetic pathway (Fig. 5*E*), was knocked out in in HAP1 cell line. As expected, MECR-null cells have no detectable level of lipoic arm on DLAT, DLST or DBT, in contrast to wildtype HAP1 cells (Fig. 5*F*). We then treated wildtype or MECR knockout cells with NO donor. The levels of functional lipoic arm on α-ketoacid dehydrogenase complexes’ E2 subunits were reduced upon NO donor treatment (Fig. 5*F*), and consistently, overall PDHC activity was reduced in wildtype HAP1 cells (Fig. 5*G*). MECR knockout cells have no PDHC activity as expected (Fig. 5*G*). Importantly, when we specifically measured the activity of DLD, we found that DLD activity was only significantly decreased after NO donor treatment in wildtype, but not MECR knockout cells (Fig. 5*H*). This result provided in-cell evidence that the inhibition of DLD by RNS depends on the lipoic arm on E2 subunit.

## Discussion

Mitochondrial α-ketoacid dehydrogenase complexes catalyze crucial reactions at the crossroads within the metabolic network. Here we demonstrate that RNS can strongly inhibit BCKDC. This study, together with our recently published work, revealed that RNS are capable of broadly inhibiting all the α-ketoacid dehydrogenase complexes through a common mechanism— modifying and inactivating the catalytic lipoic arm of their E2 subunits. We demonstrated this mechanism drives significant alterations in the metabolism of carbohydrates and amino acids across multiple cell types, including macrophages, myotubes and myoblasts, under conditions in which NO production is induced. It has been previously reported that NO production in cytokine activated muscle cells has important physiological effects, including impairment of myoblast proliferation and differentiation (39), and induction of apoptosis in aging-induced sarcopenia (40). The inhibition of α-ketoacid dehydrogenase complexes by RNS, and the resulting alterations in mitochondrial metabolism, may play a role in mediating these effects. Given that purposeful production of NO by iNOS, eNOS (primarily expressed in endothelial cells), and nNOS (primarily expressed in neurons), as well as the generation of RNS as metabolic by-products, occurs in many physiological and pathological contexts (28, 41), this mechanism is likely to have broad significance in regulating metabolism. Alterations of these enzymes by RNS have the potential to have numerous downstream impacts via mechanisms including affecting protein acetylation and succinylation by altering acetyl-CoA and succinyl-CoA availability or changing cellular energetic status. The broad downstream effects and their mechanisms remain to be examined.

The α-ketoacid dehydrogenase complexes are subject to tight regulation by a variety of mechanisms. Here we investigated the relationship between the RNS-driven inhibitory modifications on the E2 subunit and other regulatory post-translational modifications targeting α-ketoacid dehydrogenase complexes’ E3 or E1 subunits. We found that RNS-driven modification of the E2 subunit’s lipoic arm promotes inhibition of the E3 subunit. This molecular connection has important implications in the specificity of RNS-driven DLD inhibition and in the extent and reversibility of overall α-ketoacid dehydrogenase inhibition by RNS. In high RNS conditions, such as classical activation of macrophages, we previously found that α-ketoacid dehydrogenases are specifically and profoundly inhibited, while cell viability and the activity of many other mitochondrial enzymes remains high (42). If RNS cause inhibitory DLD nitrosylation by direct, non-enzymatic interaction with DLD, it is mechanistically unclear why DLD would be preferably modified and inhibited, over many other mitochondrial proteins that have cysteine residues which can potentially be modified. This specificity question is explained by our model of multi-step modification delivery: In cells, RNS can react with CoA, a relatively abundant thiol containing metabolite, and generate SNO-CoA. Through the specific binding of SNO-CoA to the E2 subunit, the modification is efficiently delivered to the thiol of lipoic arm; and through the local interaction between the E2 and E3 subunits, the E3 subunit is further modified. Through this mechanism, both E2 and E3 subunits can be inactivated by RNS, causing greater effect on overall α-ketoacid dehydrogenase inhibition. As a result, to recover the overall enzyme activity, the inhibitory modifications on both E2 and E3 subunits need to be removed. The reversibility under specific cellular conditions is an important direction for future studies.

Overall, this study extended our knowledge about the mechanisms impacting the activity of α-ketoacid dehydrogenase complexes and provided insight into the relationship among different regulatory mechanisms which act upon these enzymes. This work showed that strong inhibition of α-ketoacid dehydrogenase complexes by RNS can have significant effects in cellular metabolism across various cell types. These findings merit future investigation to examine the broader physiological or pathological effects of this mechanism *in vivo* and explore the translational implications in conditions where elevated RNS play a key role, such as inflammatory disorders and cardiovascular diseases.

## Experimental Procedures

### Cell culture

RAW 264.7 cells, wild type or *Nos2*−/−, were cultured at 37°C with 5% CO_2_ in RPMI 1640 media with 1% penicillin-streptomycin, 25 mM HEPES, and 10% fetal bovine serum (FBS). Dialyzed fetal bovine serum (dFBS) was used in the place of FBS for metabolomics and isotope tracing experiments. Media was replaced every 24 hours. To stimulate the cells, 50 ng/mL lipopolysaccharide (LPS) (E. coli 0111:B4) (Sigma-Aldrich, L3024) and 10 ng/mL recombinant mouse interferon-γ (IFN-γ) (R&D Systems, 485-MI-100) were added to the media and maintained with subsequent media change.

Murine myoblast cell line, C2C12 (ATCC), were cultured in growth media (Dulbecco’s Modified Eagles Medium (DMEM) with 10% FBS and 1% penicillin-streptomycin) at 37°C with 5% CO_2_. To differentiate C2C12 cells to myotubes, media was replaced with differentiation media (DMEM with 2% Donor Equine Serum (Hyclone) and 1% penicillin-streptomycin) once cells reached 70-90% confluency. Media was replaced every 48-hours for five days. After differentiation, media was changed to DMEM with 2% FBS, or dFBS (for metabolomic or isotopic tracing experiments), and media was refreshed every 24 hours. For cytokine stimulation, 20 ng/mL recombinant mouse tumor necrosis factor-α (TNF-α) (R&D Systems, 41-0MT0-25CF) and 12 ng/mL recombinant mouse IFN-γ were added to the media.

The human chronic myeloid leukemia haploid cell line, HAP1, wild type or MECR−/−, were cultured in Improved Modified Eagles Medium (IMEM) with 10% FBS and 1% penicillin-streptomycin at 37°C with 5% CO_2_. For NO donor treatment, 250 µM of DETA-NONOate was added to the media for 48-hours with media replacement every 24 hours.

All cell lines were tested for mycoplasma contamination.

For leucine tracing experiments, media without L-leucine was supplemented with U-[^13^C]-L-leucine (Cambridge Isotope Laboratories, CLM-2262-H) at formulation concentration and was used in the place of chemically identical regular unlabeled media. Both RAW264.7 and C2C12 cells were cultured with stable isotope for 48 hours with media changes at 24- and 2-hours prior to metabolite extraction. For kinetic glucose tracing, media without D-glucose supplemented with U-[^13^C]-D-glucose (Cambridge Isotope Laboratories, CLM-1396-1) at formulation concentration was used.

For iNOS inhibitor treatment of C2C12 myotubes, 200 µM 1400W (Cayman Chemical, 81520) was added to the media 24 hours prior to experiment start as pre-treatment. The inhibitor was maintained in the media at the same concentration throughout the experiment duration.

### CRISPR-Cas9 based genetic knockout of Nos2

*Nos2*−/− knockout cells were generated as previously described (24). Briefly, 2 x 10^6^ C2C12 myoblast cells were transfected via electroporation with 1 µM fluorescent trans-activating CRISPR RNA (tracerRNA, ITT, catalog no. ATTO550), 1 µM RNA targeting mouse *Nos2* (crRNA, GTGACGGCAAACATGACTTC, IDT Design ID: Mm.Cas9.NOS2.1.AA) and 1 µM HiFi Cas9 enzyme (IDT, catalog no. 1081060) in 100 µL Nucleofector Solution V plus supplement (Lonza, catalog no. VCA-1003), using the preprogrammed electroporation protocol B-032 on Nucleofector II/2b. Immediately, cells were plated on a 35-mm plate with DMEM media with 10% FBS without penicillin/streptomycin. Eighteen hours after transfection, cells positive for fluorescent tracrRNA were single-cell sorted by FACS (BD FACSAria III) onto a 96-well plate in DMEM media with 10% FBS and 1% penicillin/streptomycin. Single-cell colonies were expanded and subsequently screened via western blot for the lack of iNOS expression after 48-hour stimulation with TNFα and IFNγ. Further validation of positive hits (myoblasts without iNOS expression after cytokine treatment) was performed by differentiating the selected myoblast clones into myotubes followed by 48-hour treatment with TNFα and IFNγ and subsequent western blot for iNOS expression, measurement of nitrite concentration in the media using Griess Reagent System (Promega, G2930), and measurement of intracellular citrulline abundance by LCMS.

### Protein extraction, SDS-PAGE, and immunoblotting

Whole cell lysate was collected using RIPA buffer (150 mM NaCl, 1% NP-40 substitute, 50 mM Tris, 0.4 mM EDTA, 0.1% SDS, 0.5% Sodium Deoxycholate, 10% (v/v) glycerol, pH=8.0). Lysate was incubated on ice for 15 minutes and spun at 12,000 x g for 5 minutes at 4°C. Total soluble protein concentration in supernatant was determined with BCA assay (Thermo Scientific, Pierce 23225). Denatured gel was run using a Thermofisher Scientific Mini Gel Tank system with Bolt Bis-Tris 8% or 4-12% gels and Bolt MES or MOPS running buffer. Proteins were then transferred to nitrocellulose membrane using Bolt Transfer Buffer. Total protein stain was used (Li-COR, 926-11011) to visualize loading. Membranes were blocked in 5% non-fat dairy milk (NFDM) in Tris-Buffer Saline with 0.01% Tween-20 (0.01% TBS-T) for 1 hour at room temperature. Primary antibodies were diluted 1:1000 in 5% bovine serum albumin (BSA) in 0.01% TBS-T and incubated with membrane overnight at 4°C. Membranes were washed with TBS-T and placed with secondary antibody diluted 1:10,000 in 5% BSA in TBS-T for 1 hour at room temperature. Blots were imaged on Odyssey CLx Imaging System (LI-COR Biosciences).

For immunoblotting, the following antibodies were used: Anti-DLST (Abcam, ab187699), Anti-DBT (Abcam, ab151991), Anti-Lipoic Acid (Millipore Sigma, 43-769-5100UL), Anti-DLAT (Abcam, ab172617), Anti-iNOS (Cell Signaling Technology, 2982S), Anti-DLD (Abcam, 133551), Anti-alpha-tubulin (Cell Signaling Technology, 3873S), Anti-beta-actin (Cell Signaling Technology, 4967S), Anti-Myosin Heavy Chain (Abcam, ab37484), IRDye 800 CW Goat Anti-Mouse IgG Secondary (LI-COR, 926-32210), and IRDye 800 CW Goat Anti-Rabbit IgG Secondary (LI-COR, 926-32211).

### Immunoprecipitation

Wild type or *Nos2*−/− RAW264.7 cells were collected into centrifuge tubes after washing twice with D-PBS and scrapping off the culture plates. Cell suspensions were spun at 500 x g for 10 minutes at 4°C and supernatant was discarded. Protein was extracted from the cell pellets using extraction buffer (20 mM Tris HCl, pH 8.0, 150 mM NaCl, 1 mM EDTA, 1 mM EGTA, 1% NP-40 with Pierce Protease and Phosphatase inhibitors (Thermo Fisher, A32957 and A32953)) at a volume of 0.350 mL per 3.00 x 10^7^ cells. Cell mixtures were incubated on ice for 30 minutes, then centrifuged at 5,000 x g for 10 minutes at 4°C. Supernatant was transferred to a tube, and total soluble protein concentration was determined with BCA assay. Dynabeads Protein G (Thermo Fisher, 10003D) (200 µL per isolation) were washed three times with Citrate-Phosphate Buffer (470 mg Citric Acid, 920 mg Dibasic Sodium Phosphate dihydrate, pH=8.0), then incubated with 2 µL of anti-lipoic acid antibody in 200 µL D-PBS with gentle mixing for 40 minutes at room temperature and washed three times with Citrate-Phosphate Buffer. Antibody was cross-linked by washing twice with 0.2 M triethanolamine (pH=8.2) and resuspending in fresh DMP solution (20 mM dimethyl pimelimidate dihydrochloride (Fisher Scientific, P08925G) in 0.2 M triethanolamine, PH=8.2 (DOT Scientific Inc., DST23040-0.1)) and incubating for 30 minutes at room temperature. The reaction was terminated by resuspending beads in 50 mM Tris buffer (pH=7.5) and incubating for 15 minutes. Cross-linked beads were washed three times with PBS with 0.1% Tween-20. To immunoprecipitate protein containing lipoic moiety, 2 mg of whole cell lysate were mixed with cross-linked bead-Ig complex and the mixture was rotated end-over-end overnight at 4°C. Then the beads were washed three times with PBS and protein was eluted with 50 µL of glycine elution buffer (50 mM glycine, pH=2.8). To ensure complete elution, an additional 30 µL of elution buffer was added to the beads, and the eluants were pooled. The eluant was then neutralized with 1:1 (v/v) neutralization buffer (1 M Tris, pH=7.5). Total protein concentration of eluant was determined with BCA assay, then the same amount was loaded for immunoblotting for DBT.

### Metabolite extraction and Liquid chromatography-mass spectrometry (LCMS) analysis

Cells were washed three times with Dulbecco’s Phosphate Buffer Saline (D-PBS) and intracellular metabolites were extracted with cold 80:20 methanol: H_2_O (v/v, LCMS grade). Samples were dried under nitrogen gas and resuspended in LCMS-grade H_2_O.

Samples were analyzed using a Thermo Q-Exactive mass spectrometer coupled to a Vanquish Horizon UHPLC. Analytes were separated on a 100 x 2.1 mm, 1.7 µM Acquity UPLC BEH C18 Column (Waters), with a gradient of solvent A (97:3 H_2_O: methanol, 10 mM TBA, 9 mM acetate, pH 8.2) and solvent B (100% methanol) at 0.2 ml/ min flow rate. The gradient is: 0 min, 5% B; 2.5 min, 5% B; 17 min, 95% B; 21 min, 95% B; 21.5 min, 5%. Data were collected in full-scan negative mode. Setting for the ion source were: 10 aux gas flow rate, 35 sheath gas flow rate, 2 sweep gas flow rate, 3.2kV spray voltage, 320°C capillary temperature and 300°C heater temperature.

The metabolites reported were identified based on exact *m/z* and expected retention times determined with chemical standards. Data were analyzed with MAVEN (47, 48).

To quantify changes in relative metabolite levels, metabolite abundance measured by ion count in LCMS analysis were normalized to total protein content. To quantify absolute abundance for AMP, ADP, and ATP to calculate cellular energy charge, the ion count measured by LCMS was converted to molar quantity based on calibration curves obtained by running various concentrations of purified AMP, ADP, and ATP chemical standards on LCMS using the same method. Energy charge is calculated based on *Equation 1*.

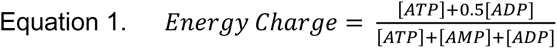

### Crude mitochondria isolation and matrix enrichment

Mitochondria isolation and matrix enrichment was performed as previously described (21, 49, 50) with adaptations. Briefly, cells were harvested from tissue culture plate after washing three times with D-PBS. Cells were pelleted by spinning at 1,000 x g for 5 minutes at 4°C. Cell pellet was resuspended in isolation buffer (10 mM Tris-MOPS, pH=7.4, 1 mM EGTA/Tris, and 200 mM sucrose) and then homogenized with Teflon pestle operating at 1,600 rpm with 50 passes. Homogenate was spun at 600 x g for 10 minutes at 4°C. Supernatant was transferred to a new tube and spun at 7,000 x g for 10 minutes at 4°C. Pellet was resuspended in isolation buffer then spun at 7,000 x g. Mitochondria pellet was lysed with hypotonic lysis buffer (20 mM Tris, pH 7.4, 1 mM EDTA, and Pierce Protease and Phosphatase Inhibitors) and incubated on ice for 15 minutes. For crude mitochondria use, mitochondria protein concentration is determined with BCA assay.

For matrix lysate enrichment, pelleted crude mitochondria were lysed with 150 µL hypotonic lysis buffer, sonicated on ice for 5 seconds followed by 30 second rest for a total of four times at 40% amplitude with a probe sonicator. After the addition of 30 µL 1 M NaCl and 20 µL 50% glycerol (v/v) to reach final concentration of 150 mM NaCl and 5% glycerol, lysate was spun at 20,000 x g for 30 minutes at 4°C. For DLD activity assays, matrix lysate was dialyzed (3.5K MWCO membrane) overnight at 4°C rocking in 20 mM sodium phosphate buffer. The protein concentration in supernatant matrix lysate was determined with BCA assay.

### Branched chain α-ketoacid dehydrogenase activity assay

BCKDC activity assay in lysate was performed as previously described with adaptations (31). To initiate reaction, 200 µL of Assay Buffer with substrate mixture (30 mM K_3_PO_4_, 2 mM MgSO_4_, 2 mM DTT (Fisher Scientific, AAJ1539706), 0.56 mM TPP (Sigma-Aldrich, C8754-1G), 0.56 mM CoA (Cayman Chemical, 16147), 1 mM NAD+ (Cayman Chemical, 16077), Triton X-100, 0.2 mM alpha-ketoisocaproate (Cayman Chemical, 21052-5), and 5 µM rotenone (VWR, 10189-314)), which was pre-warmed to 30°C, and 20 µg of crude isolated mitochondria or matrix lysate, as specified in figure legends, were mixed in each well of 96-well plate. Reaction was allowed to proceed at 30°C. At designated time points (typically every 20 minutes from reaction start to 1 hour), 50 µL or 60 µL of reaction mixture sample was collected and quenched in 4X volume (200 µL or 240 µL) of LCMS-grade methanol. Samples were spun and supernatant were dried under nitrogen gas, then resuspended in LCMS-grade H_2_O and analyzed by LCMS. The reaction rate was quantified by the production of isovaleryl-CoA over time. The slope was fitted by linear regression. As a blank control, the same amount of mitochondria lysate was mixed with assay buffer without alpha-ketoisocaproate.

For experiments involving *in vitro* treatment of mitochondria lysates with NO donor or SNO-CoA, lysate was diluted to 1.5 µg/µL in assay buffer (30 mM K_3_PO_4_, pH 7.0), PAPA-NONOate or SNO-CoA were added and incubated at room temperature for 3-5 hours, as specified in figure legend. SNO-CoA was prepared as previously described (24, 51, 52) by combining 100 mM CoA in 1 M HCl with 100 mM NaNO_2_ in 100 µM EDTA and 100 µM DPTA in a 1:1 (v/v) ratio.

### Purified Porcine PDHC Activity Assay

The PDHC activity assay was performed as previously described with adaptations (24). Briefly, purified porcine PDHC (Sigma-Aldrich, P7032-10UN) (0.7 units/mL) was incubated at room temperature for 18 hours with NO Donor, PAPA-NONOate (1 mM) (Cayman Chemical, 82410), with indicated combinations of CoA (100 µM) and pyruvate (1 mM) in 20 mM sodium phosphate buffer (pH 7.2). After incubation, the protein mixture was diluted 1:20 in assay solution containing thiamine pyrophosphate (100 µM), CoA (2 mM), pyruvate (2 mM), and NAD+ (10 mM) in 20 mM sodium phosphate buffer (pH 7.2). PDHC activity was quantified by the rate of NADH production, as measured by the increasing NADH absorbance at 340 nm over time using a BioTek Epoch2 microplate reader. Absorbance was measured continuously, and the mean velocity was determined from the linear portion of the curve. Data was analyzed using Gen5 TS v.2.09 software (BioTek Instruments, Inc.).

### Blue native-PAGE and DLD Activity Assay

For native gel analysis, the Novex Native Bis-Tris Gel System was used (Thermo Fisher). Samples were loaded in 4-16% Bis-Tris NativePAGE gel and run at 150V for 1 hour at 4°C with anode buffer in outer chamber and light blue cathode buffer in inner chamber. After 1-hour, light blue cathode buffer was replaced with anode buffer. The gel was run for an additional 1 hour at 250V on ice at 4°C.

In gel DLD activity assay was performed as previously described (21, 34, 35). Briefly, native gel was immediately removed from cassette and incubated in activity assay buffer (50 mM potassium phosphate, pH=7.0, 0.2 mg/mL nitro blue tetrazolium (NBT) chloride (Alta Aesar, B23792.02), and 0.1 mg/mL NADH (Cayman Chemical, 16078)) until purple bands appeared. The gel was then imaged on an EPSON Scan V700. After the image was obtained, gel was fixed, stained with Coomassie R-250, and de-stained for visualization of the protein standard (Thermo Fisher, LC0725). The product of the diaphorase activity of DLD is NBT-formazan, which has a maximum absorbance between 500-600nm (36). Therefore, DLD activity was also quantified by the diaphorase activity as measured by the increasing NBT-formazan absorbance at 568 nm over time using a BioTek Epoch2 microplate reader. Absorbance was measured continuously, and the mean velocity was determined from the linear portion of the curve. As a blank control, the same amount of mitochondria lysate was mixed with assay buffer without NADH. Mean velocity was normalized to total DLD protein level quantified by parallel western blot. Data was analyzed using Gen5 TS v.2.09 software (BioTek Instruments, Inc.).

### Measurement of nitrite concentration

To measure nitrite production by cells, 2 mL media was incubated with each well of cells in 6-well plate for 48 hours, then spent media was collected. Nitrite concentration in spent media was measured using Griess Reagent System (Promega, G2930) per manufacturer’s instructions.

### Statistical Analysis

The exact statistical analysis used in each experiment was stated in figure legend. In general, for comparisons between two groups, a non-paired students’ t-test was performed. For comparisons between three groups or more, one-way ANOVA followed by Tukey’s post-hoc test for multiple comparisons was performed.

### Software

LCMS data analysis was performed with Maven Version 6.2. Immunoblots were visualized and quantified using Image Studio Lite version 5.2 for Windows (LI-COR Biosciences, Lincoln, NE USA). Data were graphed, and all statistical analyses were completed in GraphPad Prism version 9.4.1 for Windows (Graph Pad Software, San Diego, California USA). Figures were created with Adobe Illustrator 2023 and ChemDraw, Professional, Version 22.0.0.22.

## Data availability

All data are contained within the article. Source data for all the figures are available upon request to the corresponding author.

## Supporting information

This article does not contain supporting information.

## Acknowledgements

FACS for sorting single colonies of *Nos2* knockout C2C12 cells was performed with the instrument and assistance of UWCCC Flow Lab (the flow cytometry core is supported by Cancer Center Support Grant P30 CA014520). This work is supported by NIH grant R35GM147014 (J.F.). We thank David J. Pagliarini (WashU) for the HAP1 cells.

## Author contributions

N.L.A., G.L.S. and J.F. conceptualized the study, designed the experiments, and analyzed the results. N.L.A and J.J. perform the experiments. J.F. supervised the study. N.L.A. and J.F. wrote the manuscript, G.L.S. edited the manuscript.

## Funding and additional information

This work is supported by NIH grant R35GM147014 (J.F.). N.L.A. is supported by the Medical Scientist Training Program (T32 GM140935).

